# Neuronal expression of S100B triggered by oligomeric Aβ peptide protects against cytoskeletal damage and synaptic loss

**DOI:** 10.1101/2025.04.04.647260

**Authors:** Joana Saavedra, Mariana Nascimento, António J. Figueira, Marina I. Oliveira da Silva, Tiago Gião, João Oliveira, Márcia A. Liz, Cláudio M. Gomes, Isabel Cardoso

## Abstract

Alzheimer’s disease (AD) is a complex neurodegenerative disorder characterized by the intracellular deposition of Tau protein and extracellular deposition of amyloid-β peptide (Aβ). AD is also characterized by neuroinflammation and synapse loss, among others. The S100 family is a group of calcium-binding proteins with intra- and extracellular functions, that are important modulators of inflammatory responses. S100B, which is upregulated in AD patients and the most abundant member of this family, was shown to inhibit *in vitro* the aggregation and toxicity of Aβ42, acting as a neuroprotective holdase-type chaperone. Although S100B is primarily produced by astrocytes, it is also expressed by various cells, including neurons. In this work, we investigated if S100B neuronal expression is triggered as a response to Aβ toxic species, to provide protection during disease progression. We used the AD mouse model AβPPswe/PS1A246E to show that neuronal S100B levels are significantly higher in 10-month-old animals, and cellular assays to demonstrate that Aβ oligomers significantly increase S100B expression in SH-SY5Y cells, but not monomeric or fibrillar Aβ. Using primary cultures of rat hippocampal neurons, we showed that S100B partially reverts Aβ-induced cofilin-actin rods (synapse disruptors), and rescues the decrease in active synapses and post- (PSD-95) synaptic marker, imposed by Aβ peptide. Altogether, these findings establish the neuroprotective activity of S100B in response to proteotoxic stress in cells, highlighting its chaperone function as a crucial factor in understanding proteostasis regulation in the diseased brain and identifying potential therapeutic targets.

## Introduction

One of the main hallmarks of Alzheimer’s Disease (AD) is the aggregation of amyloid-β peptide (Aβ) into extracellular plaques ^1–3^. In AD a complex self-assembly process converts monomeric Aβ into mature amyloid fibrils, comprising the formation of soluble/fibrillar oligomeric species, known as Aβ oligomers ^4^ that are the responsible for disease progression ^5,6^. Oligomers trigger the start of a series of harmful events that contribute to neuronal loss and cognitive decline ^7^. Examples include tau hyperphosphorylation and missorting ^8,9^, oxidative stress ^10,11^, metallostasis dysregulation ^12,13^, cell membrane damage ^14^, axonal transport impairment ^15^, synaptic receptor redistribution ^16^, astroglia activation ^17,18^ and formation of cofilin-actin rods ^19–21^. In AD, these cofilin-actin rods, structures composed of bundles of cofilin-saturated actin filaments, which result from localized cofilin hyperactivation by dephosphorylation and oxidation ^22^, inhibit intracellular trafficking and cause synaptic loss in cultured hippocampal neurons ^23,24^.

Neuroinflammation is an important characteristic commonly present in neurodegenerative disorders, and thought to play a relevant role in AD. In its acute phase, neuroinflammation is beneficial; however, persistent activation of microglia and astrocytes, such as that caused by Aβ aggregated species, is detrimental and can contribute to AD pathogenesis. The accumulation of protein aggregates and consequent increase in inflammation results in the release of inflammatory mediators, including several alarmins ^25^, such as Ca^2+^-binding S100B protein ^26,27^, which is upregulated in AD^28^ and is known to contribute to the late neuroinflammatory response ^29,30^.

The high conservation of S100 family proteins across vertebrate species underscored their essential biological roles ^31,32^. S100B is the most prevalent protein in this family, constituting 0.5% of all brain proteins and 96% of the S100 proteins found in the human brain ^33^. S100B is primarily found in brain tissue, especially in astrocytes, but is also present in other cells such as enteric glia, microglia, Schwann cells, and oligodendrocytes. It is detected at low levels in specific neuronal populations in the cerebellum, forebrain, spinal cord, and other areas. S100B has also been observed in vascular endothelial cells, lymphocytes, and various other non-neural cells, including adipocytes, chondrocytes, and melanocytes. However, it is not found in microglial cells ^33^. S100B can act intra- and extracellularly, and, depending on its concentration and the developmental stage at which it is expressed, the protein assumes distinct roles. By modulating calcium signaling and interacting with a variety of molecules in multiple cell types, S100B takes part in a number of biological activities. For instance, S100B generally controls cellular calcium homeostasis and enzyme activity, interacts with the cytoskeleton, and has a role in cell proliferation, survival, and differentiation processes ^34^. Additionally, S100B is essential to maintain blood-brain barrier functional integrity ^35^ and modulates synaptic plasticity and hippocampus-dependent memory ^36–38^. Moreover, in addition to its pro-inflammatory function, S100B acts as a neuroprotective holdase type chaperone by preventing the *in vitro* aggregation and toxicity of the Aβ42^39–41^ and of the microtubule-associated protein tau ^42^. S100B protects cells from Aβ42-mediated toxicity, rescuing cell viability and decreasing apoptosis induced by Aβ42 in cell cultures ^39^, but the mechanisms underlying this protection remain unknown. In this work, we address this aspect by testing the hypothesis that Aβ toxic species trigger neuronal S100B expression to mitigate noxious effects produced by proteotoxic Aβ over the cytoskeleton and synaptic structures.

## Material and methods

### Proteins and peptides

Synthetic Aβ42 (GenScript) was dissolved in hexafluoro-2-propanol (HFIP) (Merck) and left at room temperature (RT) for 48 hours. Subsequently, the HFIP was removed using a stream of nitrogen, and the resulting powder was dissolved in dimethyl sulfoxide (DMSO) (Sigma-Aldrich) at a concentration of 2000 µM. For oligomer and fibril formation, Aβ42 was diluted in F-12 medium (Alfagene) to a concentration of 100 µM and incubated for 48 hours at 4 ºC, and for 72 hours at 37ºC, respectively. Soluble Aβ42 was obtained by diluting, at the time of use of the peptide, to the desired concentration. Human S100B was expressed in E. coli (BL21 (DE3) E. Cline Express, Lucigen) and purified to homogeneity as described ^43,44^. S100B concentrations were estimated as homodimer equivalents using by UV spectroscopy at 280 nm using the theoretical extinction coefficient value of ε_280 nm_ = 2,980 M^-1^cm^-1^.

### Transmission Electron Microscopy

For the examination of the structure and morphology of samples containing the different Aβ42 species, 5 µL of the samples were adsorbed into carbon-coated collodion film supported on 300-mesh copper grids (Electron Microscopy Sciences, PA, USA) and negatively stained twice with 1% (m/v) uranyl acetate (Electron Microscopy Sciences, PA, USA). Grids were visualized with a JEOL (Tokyo, Japan) JEM-1400 transmission electron microscope, operated at 80kV, equipped with an Orious (CA, USA) Sc1000 digital camera, and exhaustively observed.

### Animals

Animals were housed in a controlled environment (12-hour light/dark cycles, temperature between 22 and 24 °C, humidity between 45 and 65%, and 15–20 air changes/hour), with freely available food and water. The AD mouse model AβPPswe/PS1A246E was maintained on a B6/C3H/SV129 genetic background. In this study, a total of 15 animals were used, divided into three age groups: 3 months, 7 months, and 10 months (n = 4-6 per group). Time pregnant wild-type female Wistar rats (E18; n=3) were used for the dissociated hippocampal neuronal cultures. All the experiments were approved by the Institute for Research and Innovation in Health Sciences (i3S) Animal Ethics Committee and in agreement with the animal ethics regulation from Directive 2010/63/EU.

### Tissue processing

Animals were anesthetized with a mixture of ketamine (75 mg/kg) and medetomidine (1 mg/kg) administrated by intraperitoneal injection, and perfused for 5 minutes with 40 ml of PBS. Brains were removed and fixed for 24 h at 4°C in 10% neutral buffered formalin and then transferred to a 30% sucrose solution for cryoprotection before cryostat sectioning and immunohistochemical analyses.

### S100B and βIII tubulin immunohistochemistry

Free-floating 30 μm-thick coronal mice brain sections, obtained with the cryostat, were permeabilized with 1% Triton X-100 in PBS for 15 minutes, at RT. After, they were washed with PBS (3x, 5 minutes), and incubated with 0.2 M NH_4_Cl in PBS for 15 minutes, at RT. Next, the tissues were once again washed with PBS and blocked with 1% BSA, 1% Triton X-100 in PBS for 1 hour, at RT, followed by an incubation with primary mouse anti-βIII tubulin antibody (ab7751; 1:500; Abcam) and with primary rabbit anti-S100B antibody (ab52642; 1:250; Abcam), in blocking solution, for 72 hours at 4ºC. After this incubation, tissues were washed with PBS (3x, 5 minutes) and incubated with Alexa Fluor-488 goat anti-mouse IgG antibody (A11029; 1:1500; Invitrogen) and Alexa Fluor-568 goat anti-rabbit IgG antibody (A11011; 1:1500; Invitrogen), ON at 4ºC. On the next day, tissues were washed with PBS (3x, 5 minutes) and incubated with DAPI (1:40000; Invitrogen) for 15 minutes at RT. Then, tissues were washed once again with PBS (3x, 5 minutes). All of the previous steps were performed with agitation. The brain sections were then mounted on normafrost slides silane pre-coated (Normax), and incubated with glycerol 90% in PBS (VWR), without agitation, for 45 minutes at RT. For the evaluation of colocalization between S100B and βIII tubulin, the tissues were visualized and imaged using a Zeiss Axio Imager Z1 microscope equipped with an Axiocam MR3.0 camera and Axivision 4.9.1 software. Images of the hippocampus were taken using a 40x magnification objective.

### Primary cultures of hippocampal neurons

The hippocampus was dissected from E18 rat embryos (Wistar) digested with 0.06% trypsin (Sigma-Aldrich, T4799) in Hanks’ balanced salt solution (HBSS, Sigma, H9394) for 15 minutes at 37 °C. Following digestion, neurons were dissociated by gentle trituration and resuspended in neurobasal medium (Invitrogen, 21103049) supplemented with 10% Fetal bovine serum (FBS), 2% N21-MAX (R&D Systems, AR008), 1% Pen-Strep and 2.5 mM L-Glutamine (Lonza, 17-605E). Cells were then counted and plated at a density of 20000 cells/coverslip in a 24-well plate for immunostaining analysis. Coverslips (13mm; glass; 1,5 mm thickness (Avantor)) were precoated with 20 µg/mL poly-D-lysine (Sigma, P0899). After 4h, the medium was changed to complete medium without FBS (2% N21, 1% Pen-Strep, 2.5mM L-Glu in Neurobasal medium).

### Cofilin and βIII tubulin immunocytochemistry in primary rat neurons

DIV5 rat neurons were incubated with 10 μM Aβ42 oligomers for 48 hours or at DIV6 with 5 μM S100B for 24 hours; Alternatively, S100B was added halfway through the 48-hour Aβ42 treatment. Neurons were fixed at DIV7 with 4% PFA in 1x cytoskeleton preservation buffer (10mM MES pH 6.1; 138mM KCl; 3mM MgCl_2_; 2mM EGTA pH 7; 0.32M sucrose), for 30 minutes at RT, and then rinsed 3x times with PBS, and incubated with -20 ºC methanol, for 3 minutes at RT. Next, cells were again rinsed 3x with PBS and incubated with blocking solution (2,5% donkey serum in PBS containing 1% BSA (Bovine serum albumin), for 1 hour at RT. This was followed by incubation with primary rabbit anti-cofilin antibody (D3F9; 1:2000; Cell Signaling) and mouse anti-βIII tubulin antibody (G712A; 1:2000; Promega), diluted in 1% BSA in PBS, overnight (ON) at 4ºC. On the next day, neurons were rinsed 3x with PBS and incubated with Alexa Fluor-568 goat anti-rabbit IgG antibody (A11011; 1:1000; Invitrogen) and Alexa Fluor-488 goat anti-mouse IgG antibody (A11029;1:1000; Invitrogen), in 1% BSA in PBS, for 1 hour at RT. After incubation, neurons were rinsed 3x with PBS and coverslips were mounted with FluoroshieldTM with DAPI (Sigma-Aldrich). Cell visualization and image capture was done using the Zeiss Axio Imager Z1 microscope equipped with an Axiocam MR3.0 camera and Axivision 4.9.1 software, with a 40x magnification. Finally, cells with cofilin-actin rods were manually counted and divided per total number of cells.

### VGLUT1/PSD-95 immunocytochemistry in primary rat neurons

DIV12 rat neurons were incubated with 10 μM Aβ42 oligomers for 48 hours or at DIV13 with 5 μM S100B for 24 hours; Alternatively, S100B was added halfway through the 48-hour Aβ42 treatment. Neurons were fixed at DIV 14 with 2% PFA in PBS for 3 minutes, on ice, and then washed with ice cold PBS, and incubated with methanol for 10 minutes, at -20 ºC. Next neurons were rinsed 3x with PBS and incubated in 0.2 M NH_4_Cl in dH_2_O, for 5 minutes, at RT, followed by another rinse 3x with PBS. Then, cells were blocked with 2% BSA, 2% FBS, 0,2% fish gelatin in PBS, for 30 minutes at RT. After, neurons were incubated with primary rabbit anti-VGLUT1 antibody (135B303; 1:1000; Synaptic Systems), mouse anti-PSD-95 antibody (MA1-045; 1:1000; Thermo Fisher Scientific), and guinea pig anti-βIII tubulin antibody (302404 ;1:2500; Synaptic Systems), in blocking buffer (diluted 10x), ON at 4 ºC. On the next day, neurons were rinsed 3x with PBS and incubated with the secondary antibodies Alexa Fluor-568 donkey anti-rabbit IgG antibody (A11011; 1:1000; Invitrogen), Alexa Fluor-647 donkey anti-mouse IgG antibody (A11029; 1:1000; Invitrogen), Alexa Fluor-488 donkey anti-guinea pig IgG antibody (706-545-148; 1:1000; Jackson ImmunoResearch), in blocking buffer (diluted 10x), for 30 minutes at RT. After this incubation, cells were rinsed 3 times with PBS and coverslips were mounted with FluoroshieldTM with DAPI (Sigma-Aldrich). Visualization of cells and image capture was done using the Zeiss Axio Imager Z1 microscope equipped with an Axiocam MR3.0 camera and Axivision 4.9.1 software, with a 40x magnification. VGLUT1 and PSD-95, pre-and post-synaptic markers, respectively, were also used to the analysis of the active synapses, by assessing the colocalization of both markers. In this experiment, only one neurite per neuron was analyzed and the number of VGLUT1, PSD-95 and active synapses per neuritic length was quantified using Fiji software ^45^. An average of 30 neurons was quantified in each condition.

### SH-SY5Y cell line

The SH-SY5Y cell line (originated from the SK-N-SH neuroblastoma cell line) was used between passage 25 and 30, and cultured in Dulbecco’s Modified Eagle Medium (DMEM) F12 (Alfagene) supplemented with 10% FBS (Gibco) and 1% Pen-Strep and cells were incubated at 37°C in a humidified atmosphere with 5% of CO_2_. The medium was changed every 2-3 days.

### Cells treatments with Aβ42

SH-SY5Y cells were studied under different conditions including incubation with 10 μM Aβ42 soluble, oligomeric or fibrillar, different concentrations of oligomeric Aβ42 (1-10 μM), and for different time periods (12, 24 and 48 hours). After incubation, cells were analysed by real time-qPCR, immunocytochemistry, while supernatants were investigated by slot blot.

### Real Time-qPCR

SH-SY5Y cells were homogenized with trizol reagent (Ambion) for 5 minutes at RT, and transferred to an eppendorf. Then, chloroform (Sigma) was added (0.2 mL of chloroform for each 1 mL of trizol), and the eppendorfs were shaken for 15 seconds, followed by a 2-3 minutes incubation at RT. After centrifugation at 12000 g, for 15 minutes, at 4ºC, the upper phase was collected in a new eppendorf. Isopropanol (Sigma) (0.5mL of isopropanol for each 1mL of trizol) was added, followed by a 10 minutes incubation at RT. After another centrifugation at 14000 g for 20 minutes at 4°C, the supernatant was discarded, and the pellet was resuspended in 75% ethanol (1mL of ethanol for each 1 mL of trizol), followed by centrifugation at 7500 g for 5 minutes at 4 ºC. The supernatant was once again discarded and the ethanol was allowed to completely evaporate. The pellet was resuspended in 15-20 µL of DEPC-Water (Fisher Scientific) and left for 10 minutes at 60°C.

RNA concentration and purity were determined using a NanoDrop photometer (Thermo Fisher Scientific) by measuring the absorbance at 260 nm for RNA concentration, and the A260/A280 ratio for RNA purity assessment.

Subsequently, 2 μg of RNA were first-reversed transcribed into cDNA using NZY First-Strand cDNA Synthesis kit (ZNYtech) and the resulting cDNA was saved at -20 ºC until used. The analysis of the expression of the S100B gene was conducted using Real-Time Polymerase Chain Reaction (RT-qPCR). The actin gene was used as an internal control for normalization. The RT-qPCR reaction mixture was prepared with the SYBR green reporter (iQ SYBR green supermix, BioRad), following the manufacturer’s instructions. The PCR primers used were hActin (Forward: 5’ACAGAGCCTCGCCTTTGCCG; Reverse: 5’CACCATCACGCCCTGGTGCC) and hS100B (Forward: 5’AGAGCAGGAGGTTCTGGA; Reverse: 5’TCGTGGCAGGCAGTAGTA) and conditions used were initial denaturation 95 ºC 3 minutes, denaturation and annealing (40 cycles) 95 ºC during 10 seconds and 60 ºC during 30 seconds, melt curve 65 ºC to 95 ºC (increment 0,5 ºC for 5 seconds). The RT-qPCR was carried out using the CFX96 Touch Real-Time PCR Detection System (BioRad). The relative quantification was performed using the comparative method (2-ΔΔct) ^46^.

### Slot blot

To perform the Slot-blot analysis, 1 mL of SH-SY5Y culture supernatant was used and applied to the slot-blot manifold. The samples were drawn by vacuum through a pre-wetted nitrocellulose membrane (0,2 µm thickness) (GE Healthcare Life Science). Then, the membrane was blocked for 1h with 5% milk in PBS, at RT, and then incubated with the primary rabbit anti-S100B antibody (1:1000; Abcam) in 3% milk, ON at 4ºC. On the next day, the membrane was washed with PBS-T (3x, 5 minutes) and incubated with the secondary rabbit antibody HRP (1:2500; Binding Site), for 1h at RT. After washing again the membrane with PBS-T (3x, 5 minutes), the blot was developed using ClarityTMWestern ECL substrate (Bio-Rad), and S100B was detected and visualized using a chemiluminescence detection system (ChemiDoc, Bio-Rad).

### S100B immunocytochemistry and staining with Phalloidin iFluor-488

After fixation SH-SY5Y cells were washed with PBS (3x, 5 minutes) and permeabilized with 0,1% Triton X-100 in PBS for 10 minutes at RT. Cells were once again washed with PBS (3x, 5 minutes) and blocked with 5% BSA in PBS for 1 hour, then incubated with primary rabbit anti-S100B antibody (ab52642; 1:200; Abcam) in 1% BSA in PBS, ON at 4 ºC. On the next day, cells were washed with PBS (3x, 5 minutes) and incubated with Alexa Fluor-568 goat anti-rabbit IgG antibody (A11011; 1:1500; Invitrogen) with 1% BSA in PBS for 1 hour at RT, and then washed once again with PBS (3x, 5 minutes) and incubated with Phalloidin iFluor-488 (1:100; Abcam) in 1% BSA in PBS, for 1 hour at RT. After this incubation cells were rinsed with PBS and coverslips were mounted with FluoroshieldTM with DAPI (Sigma-Aldrich). Cell visualization and image capture were done using the Zeiss Axio Imager Z1 microscope equipped with an Axiocam MR3.0 camera and Axivision 4.9.1 software, with a 40x magnification. Fluorescence mean intensity per area in the images was quantified using Fiji software ^45^.

### Statistics

All quantitative data were expressed as mean ± standard deviation (SD). Initially, data was assessed whether it followed a Gaussian distribution. The Shapiro-Wilk test was used to assess the normality of the data and established that all data followed a normal distribution. Differences among conditions or groups were analyzed by one-way ANOVA with the appropriate post hoc pairwise tests for multiple comparisons tests. P-values lower than 0.05 were considered statistically significant. Statistical analyses were carried out using GraphPad Prism 8 software for Windows.

## Results

### Neuronal expression of S100B in the cortex increases with age

To evaluate the variation of S100B levels along the AD continuum we resorted to the AD mouse model AβPPswe/PS1A246E bearing two AD-related transgenes (APP and PSEN1) and in which Aβ deposition starts at about the age of 6 months ^47^. To investigate this, we analyzed S100B expression in animals aged 3, 7, and 10 months, representing pre-symptomatic, early amyloid deposition, and advanced disease stages, respectively. The results showed a progressive increase in S100B expression in the cortex with age, reaching significantly higher levels at 10 months compared to 3 months (Figure 1A). In contrast, no differences were observed in the hippocampus (Figure 1A). These findings align with the documented elevation of S100B levels in advanced stages of AD.

**Figure 1.**
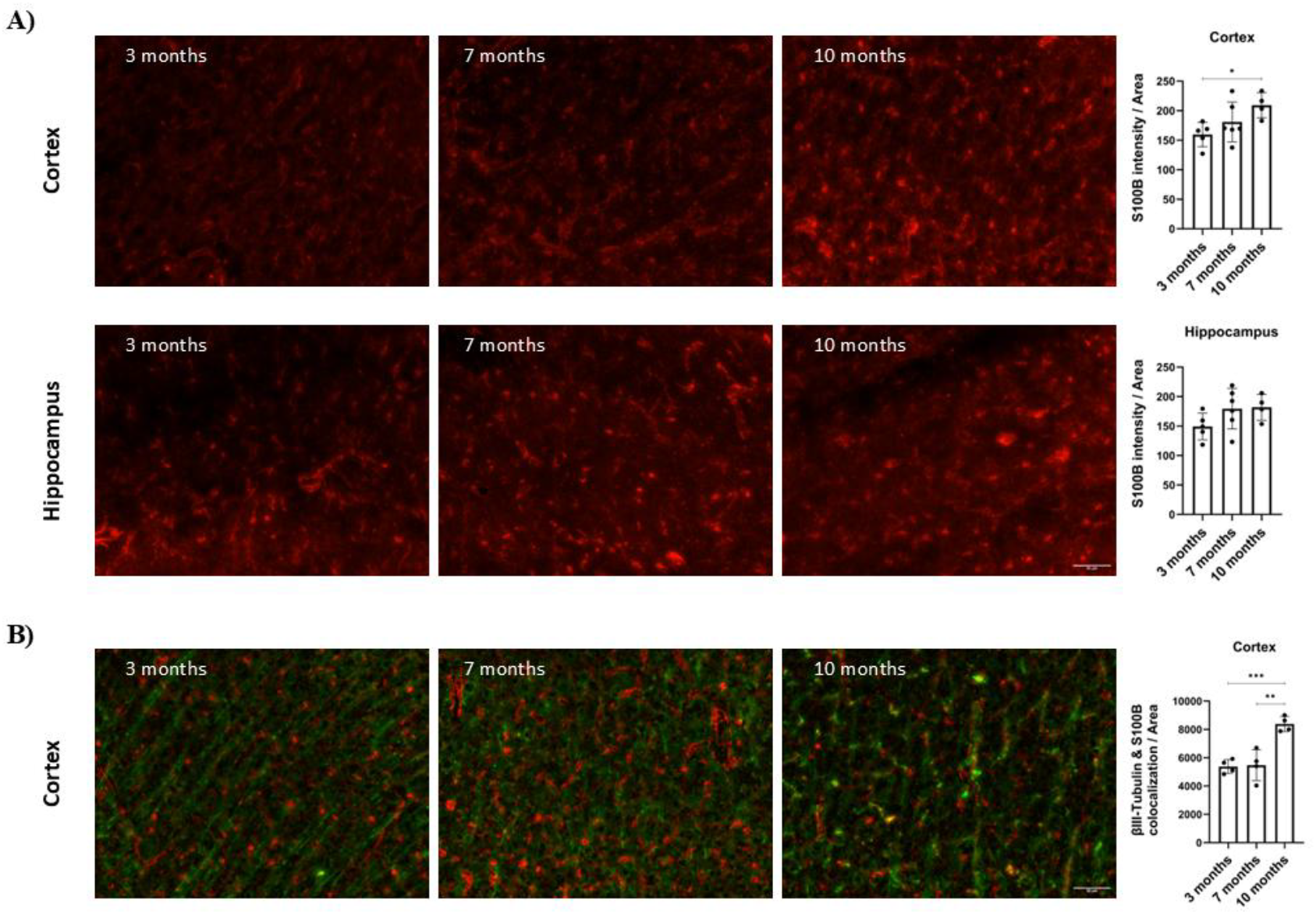
Expression of S100B in the AβPPswe/PS1A246E AD mouse model at 3, 7 and 10 months. **(A)** Representative images and quantification plots of S100B expression in AD mice at 3 (n=5), 7 (n=6) and 10 (n=4) months, showing significant differences between 10-month-old animals when compared with 3-month-old animals in the cortex. **(B)** Representative images and quantification plot of S100B colocalization with βIII-tubulin showing a significantly increase in 10-month-old animals (n=4) when compared with 3-(n=4) and 7-(n=4) month-old animals. Red: S100B; Green: βIII-tubulin. Scale bar = 50 µm. Data are expressed as mean ± SD.*p<0.05; **p<0.01; ***p<0.001.

Although S100B is a widely recognized astrocytic protein, some studies also report its expression in neurons^30^, albeit at lower levels. Given that S100B suppresses both Aβ and Tau aggregation^39,42^, we hypothesized that its neuronal expression could partially contribute to the observed protective effect. To investigate this, we analyzed cortical neuronal expression of S100B by assessing its colocalization with βIII-tubulin, a neuronal marker. The results revealed a significant increase in neuronal S100B expression in 10-month-old animals compared to both 3- and 7-month-old animals (Figure 1B), suggesting that neuronal S100B expression indeed rises with disease progression.

### Aβ oligomers induce an increase in S100B levels in the SH-SY5Y cell line

To investigate whether S100B neuronal expression is a protective response to the elevated Aβ levels observed in these animals, we conducted experiments using the SH-SY5Y cell line. Cells were treated with different Aβ species (10 μM, soluble, oligomers and fibrils) (Figure 2A), and only Aβ oligomers were found to trigger S100B expression, both at protein (Figure 2B) and transcript levels (Figure 2C). Subsequently, we tested varying concentrations of Aβ oligomers (1, 2.5, 5 and 10 μM) for 24 hours, with 10 μM Aβ oligomers significantly increasing S100B expression compared to control cells (Figure 2 D). To further refine these observations, we examined the effect of incubation duration with 10 μM Aβ oligomers (0, 12, 24 and 48 hours) finding that S100B expression peaked at 24 hours (Figure 2E). Additionally, we evaluated whether Aβ oligomers influence S100B secretion by analyzing the supernatants of cells incubated with 10 μM Aβ oligomers for 24 hours. The results revealed increased levels of secreted S100B (Figure 2F), corroborating the role of Aβ oligomers in upregulating S100B expression. Altogether, these findings suggest that neuronal S100B expression may serve as a response to Aβ oligomers.

**Figure 2.**
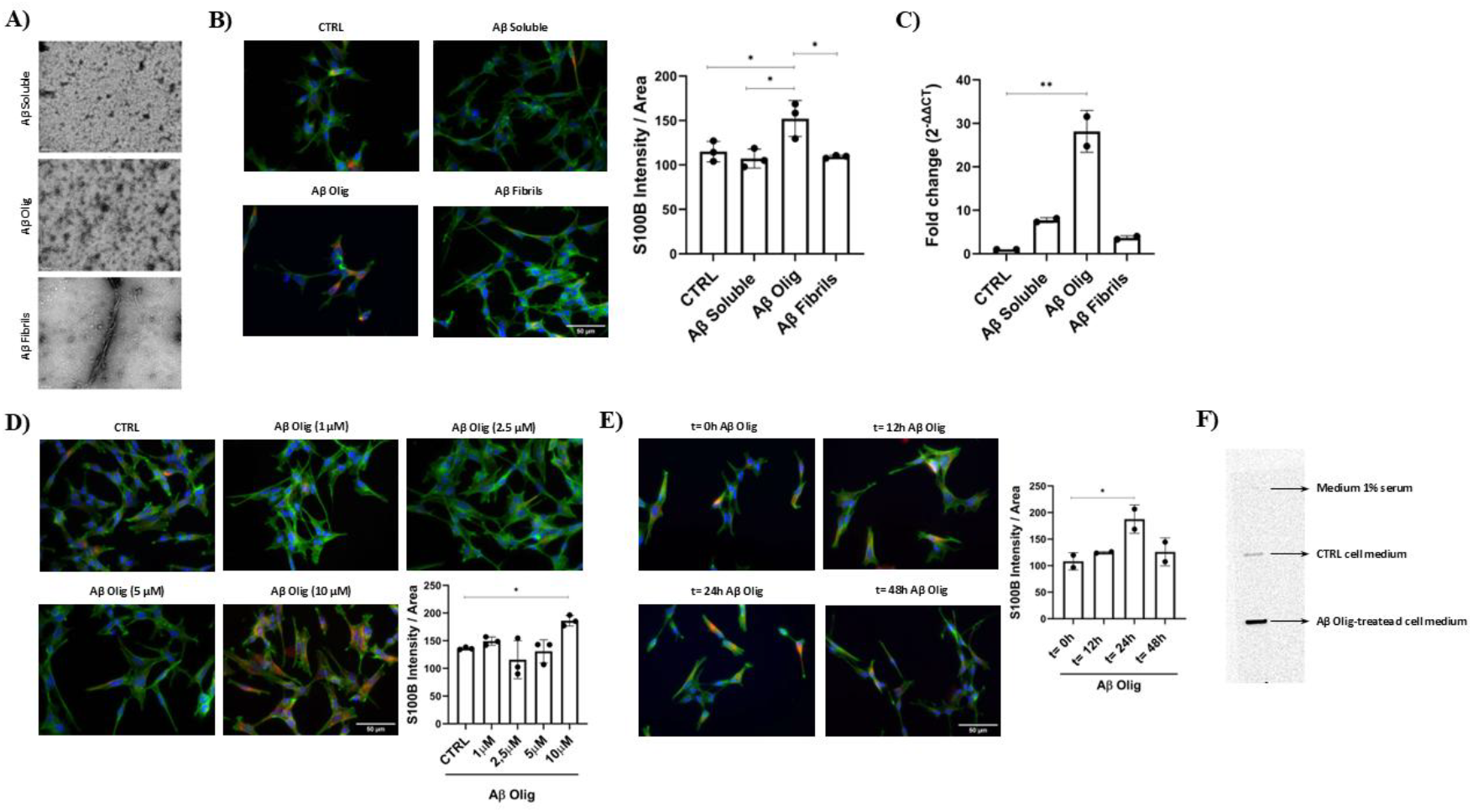
Impact of Aβ oligomers in S100B expression in SH-SY5Y cells. **(A)** TEM images of the different Aβ species (soluble, oligomeric (AβOlig) and fibrillar) used in SH-SY5Y treatments. Representative images and quantification plots of cells treated with 10 μM Aβ species (soluble, oligomeric and fibrillar), displaying different effects regarding S100B expression, with only oligomers increased S100B, both **(B)** protein (n=3 per group) and **(C)** transcript (n=2 per group). Representative images and quantification plots of **(D)** cells treated with different concentrations (1, 2.5, 5, 10 µM) of Aβ oligomers for 24 hours (n=3 per group), showing a significant increase of S100B in cells treated with Aβ oligomers at 10 µM, and **(E)** cells incubated with Aβ oligomers 10 µM for different periods of time, evidencing increased S100B in cells incubated for 24 hours, as compared to 0, 12 and 48 hours (n=2 per group). (**F**) 24 h-conditioned media, analyzed by slot blot, from cells control (Control cell medium), treated with 10 μM Aβ oligomers (Aβ Olig-treated cell medium), showing that Aβ oligomers induced increased secretion of S100B into the media. Fresh medium was used as negative control (medium 1% serum). Blue: DAPI; Green: Phalloidin; Red: S100B. Cells Scale bar = 50 µm. TEM Scale bar = 100 nm. Data are expressed as mean ± SD.*p<0.05; **p<0.01.

### S100B reverts cofilin-actin rod formation induced by Aβ42 oligomers in rat primary hippocampal neurons, protecting against cytoskeleton damage

Cofilin-actin rods are cytoskeleton inclusions that form in response to toxic stimuli, such as chemical or physical stress. These protein inclusions are often associated with neurodegenerative diseases, and their formation has been observed not only *in vitro* ^48^, but also in human AD brains ^49^ as well as in the AD mouse model used in this work (Supplementary Figure S1). Their formation disrupts normal neuronal cytoskeleton function, leading to compromised neuronal processes and overall cellular dysfunction. To further explore whether S100B, which is either expressed or internalized by neurons (Supplementary Figure S2), can mitigate toxic effects of Aβ, primary rat hippocampal neuron cultures were used to assess cofilin-actin rods formation. Neurons were exposed to Aβ oligomers (10 μM) for 48 hours to induce rod formation, with S100B (5 μM) added at the 24-hour mark to evaluate its protective effects. The results demonstrated that S100B could partially rescue the Aβ-induced phenotype, while S100B alone had no impact on rod formation (Figure 3). The protective effect of S100B displayed a dose-dependent response. Furthermore, the comparable number of cells with rods induced by Aβ at 24 and 48 hours (Supplementary Figure S3) suggests that S100B reverses rod formation rather than preventing it. These findings indicate that S100B protects against early cytoskeleton alterations triggered by Aβ oligomers, potentially mitigating disease progression.

**Figure 3.**
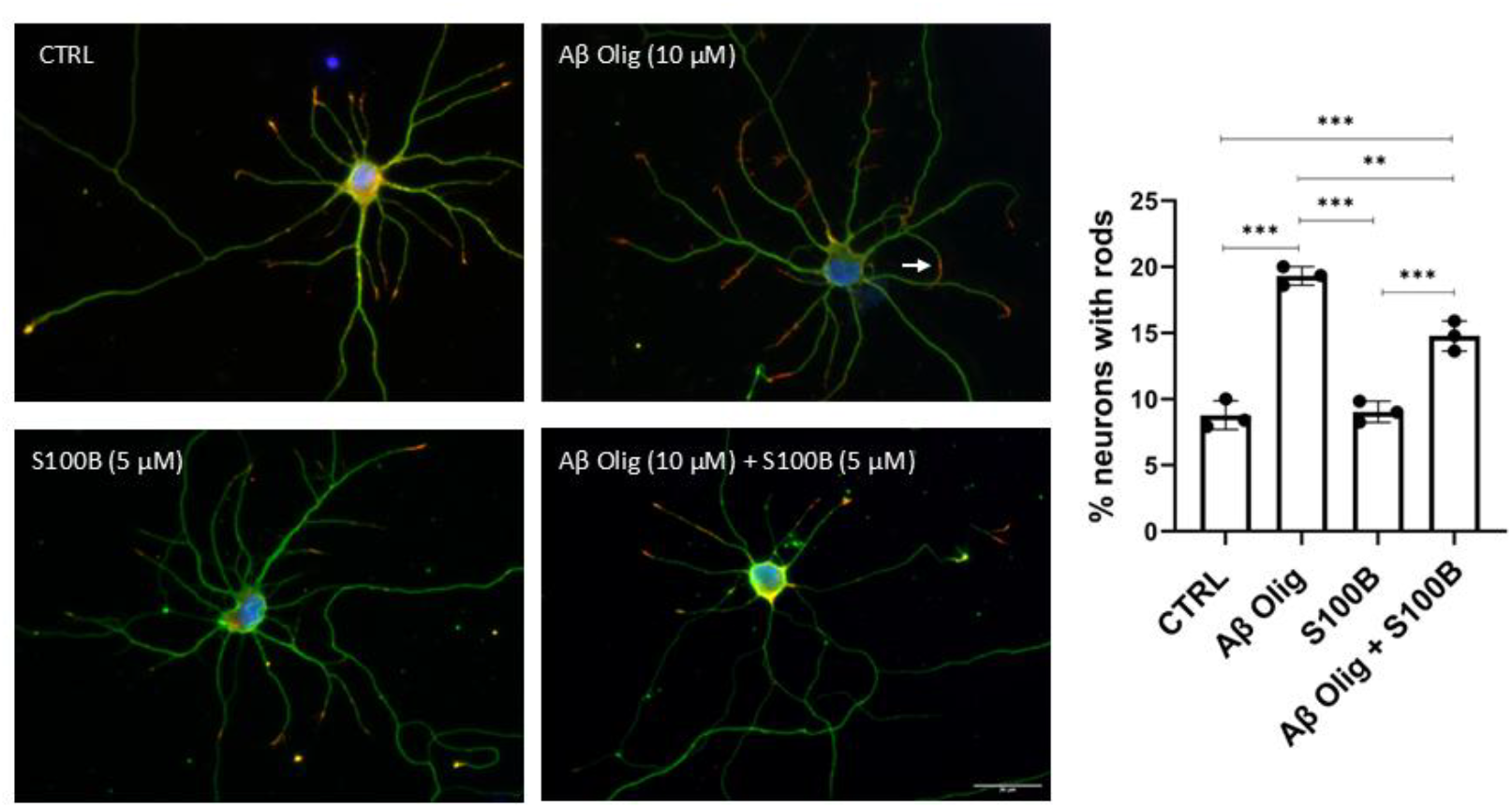
Effect of Aβ oligomers and impact of S100B in cofilin-actin rod formation in primary cultures of rat hippocampal neurons. Representative images and quantification plots of rat hippocampal neurons (DIV7) showing increased % of cells with cofilin-actin rods after incubation with Aβ oligomers (Aβ Olig, 10 μM). S100B (5 μM) added after 24 hours partially rescued the Aβ-induced phenotype. S100B alone did not produce alterations, as compared to control (CTRL). White arrow points out one cofilin-actin rod. Green: βIII-Tubulin; Red: Cofilin. Blue: DAPI. Scale bar = 30 μm. Data are expressed as mean ± SD. (n=3 per group). **p<0.01; ***p<0.001.

### S100B prevents synapse loss induced by Aβ42 oligomers in rat primary hippocampal neuron cultures

Aβ deposition in the brain leads to synaptic dysfunction by impairing neurotransmitter release and reuptake, reducing synaptic density and by triggering the formation of cofilin-actin rods, structures that inhibit intracellular trafficking and cause synaptic loss. To evaluate whether S100B directly impacts synaptic structures and can counteract the Aβ-induced dysfunction, primary cultures of rat hippocampal neurons were analyzed using immunocytochemistry for pre-and post-synaptic markers, VGLUT1 and PSD-95, respectively. As shown in Figure 4, treatment with Aβ oligomers (10 μM, 48 hours) significantly reduced PSD-95 levels and the number of active synapses compared to the control. Remarkably, S100B treatment (5 μM, 24 hours) rescued the Aβ-induced phenotype. S100B alone did not produce alterations, as compared to the control. These findings show that extracellular S100B mitigates Aβ-induced synaptic toxicity, consistent with its role as a molecular chaperone.

**Figure 4.**
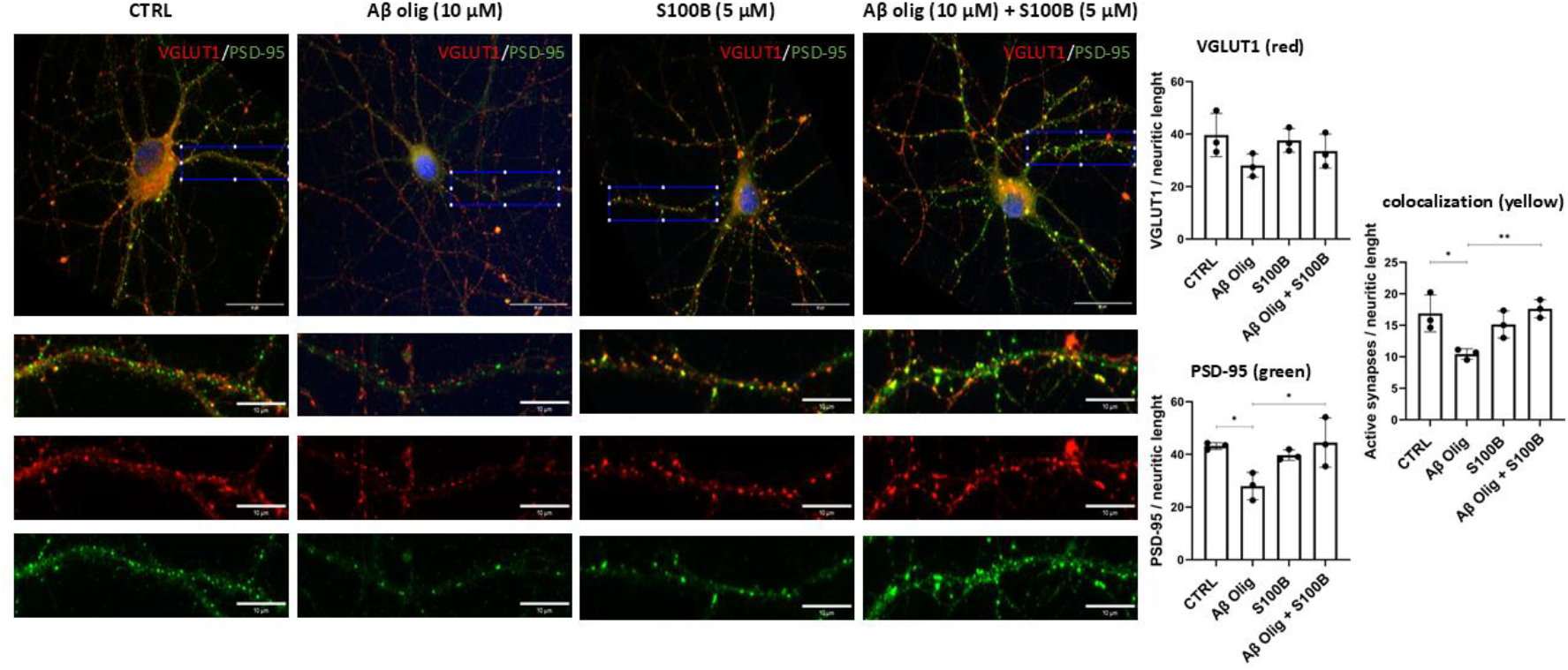
Impact of S100B and Aβ oligomers in synaptic structures of primary cultures of rat hippocampal neurons. Representative images and quantification plots of rat hippocampal neurons (DIV14) showing pre-(VGLUT1) and pos-(PSD-95) synaptic markers staining, and active synapses assessed by the colocalization of VGLUT1 and PSD-95; Treatment with Aβ oligomers (Aβ Olig, 10 μM) induces a significant decrease in PSD-95 and active synapses, as compared to control, and S100B (5 μM) rescues the Aβ-induced phenotype. S100B alone did not produce alterations, as compared to control (CTRL). Blue: DAPI; Red: VGLUT1; Green: PSD-95; Enlarged insertion (scale bar = 10 μm) corresponds to the area defined by the blue rectangle in the top panels (scale bar = 30 μm) and was used to the quantification. Data are expressed as mean ± SD. (n=3 per group). *p<0.05; **p<0.01.

## Discussion

AD is characterized by the extracellular deposition of aggregated Aβ peptide and the intracellular accumulation of Tau ^50,51^. Recent research has demonstrated that S100B, which has both intra and extracellular functions, can inhibit both the aggregation and toxicity of these proteins, even under sub-stoichiometric conditions ^39–41^. The interaction between S100B and Aβ not only reverses the peptide’s toxicity, which otherwise decreases cell viability and promotes apoptosis ^39^, but also prevents Tau oligomers from seeding further aggregation into nearby cells ^42^. These findings suggest that S100B acts in a chaperone-like capacity, mitigating the pathological effects of Aβ and Tau in AD. However, the molecular and cellular mechanisms underlying S100B neuroprotective effects remain incompletely understood, particularly those involved in neuronal protection. In this study, we demonstrate that in AD mice S100B levels increase with age, not only in astrocytes but also in neurons. This suggests that neuronal S100B expression may play a role in counteracting the toxic effects of aggregating proteins, although we cannot rule out the possibility that part of the S100B detected in neurons is partially of astrocytic origin. To further investigate this, we examined various Aβ conditions, including different species, concentrations, and incubation times, and found that only oligomers at 10 μM and incubated for 24 hours significantly increased S100B expression in SH-SY5Y cells. Notably, these neuronal cells also secreted S100B into the medium, indicating that S100B-producing neurons may help protect neighboring neurons.

The AβPPswe/PS1A246E AD mouse model employed in this study presents cofilin-actin rod formation, as observed in the brains of AD patients ^49^. Cofilin is a protein that binds actin, modulating its transport and functioning. Toxic stimuli, such as Aβ, disrupts this interaction, leading to the formation of cofilin-actin rods. These structures cause cytoskeletal dysregulation and synaptic impairment, ultimately compromising cell viability ^52,53^. Using primary rat hippocampal neuron cultures treated with Aβ oligomers, we recapitulated cofilin pathology and demonstrated that S100B partially reversed rod formation, thereby helping to preserve cytoskeletal integrity.

Another consequence of Aβ peptide deposition is synaptic dysfunction, which disrupts neurotransmitter release and reuptake, leading to reduced synaptic density. Studies in Wistar rats have shown that an intracerebroventricular injection of Aβ oligomers results in a decrease in synaptic density, structure and activity ^54^. Aβ can also block axonal transport ^55^, impair synaptic vesicle dynamics ^56,57^, and interfere with receptors in the synaptic space. Additionally, Aβ-induced dystrophic axons are linked to disruptions in the pre-synaptic machinery ^58–61^, further inhibiting neurotransmitter release and impairing synaptic function. To assess the impact of extracellular S100B in synapses, we evaluated pre- and post-synaptic markers and their co-localization to measure active synapses. Our results showed that S100B protected against the deleterious effects of Aβ, while S100B alone did not affect the synaptic structures, suggesting that its protective effect is specifically related to its ability to mitigate Aβ toxicity. In this context, cytoskeleton changes triggered by Aβ lead to the formation of cofilin-actin rods, causing synaptic damage. However, S100B, through its interaction with Aβ, prevents this sequence of events and protects synaptic integrity.

In summary, our study presents compelling evidence supporting the neuroprotective role of S100B, demonstrated both in cells *in vitro* and in the AβPPswe/PS1A246E AD mouse model. We reveal that S100B safeguards neurons by preventing the formation of cytoskeletal-disrupting cofilin-actin rods and protecting synaptic structures. These findings underscore the critical role of S100B in neuronal pathophysiology, and further identification of S100B-associated mechanisms might enable the development of therapeutic targets in AD.

## Supporting information

Supplementary material_Saavedra et al

## Supporting information

This article contains supporting information.

## Acknowledgements

The authors acknowledge the support of the i3S Scientific Platforms, Advanced Light Microcopy (ALM), member of the national infrastructure PPBI - Portuguese Platform of Bioimaging (PPBI-POCI-01-0145-FEDER-022122), Histology and Electron Microscopy platform, Cell Culture and Genotyping platform and the i3S Animal Facility.

## Author contributions

IC and CG conceived, designed and supervised the study. JS, IC, MAL and MIOS designed the experiments. JS and MN performed the experimental work and analyzed the data. AF produced the S100B protein. JS and TG collected the animals’ brains and processed them. JO cut the brains in the cryostat. IC, CG and JS wrote the manuscript. All authors revised and approved the manuscript.

## Availability of data and materials

The datasets used and/or analysed during the current study are available from the corresponding author on reasonable request.

## List of Abbreviations

AD: Alzheimer’s disease
Aβ: amyloid-b peptide
DMEM: Dulbecco’s Modified Eagle Medium
FBS: Fetal bovine serum
HBSS: Hanks’ balanced salt solution HBSS
HFIP: hexafluoro-2-propanol
ON: overnight
RT: room temperature

