## Supplementary material_Saavedra et al for "Neuronal expression of S100B triggered by oligomeric Aβ peptide protects against cytoskeletal damage and synaptic loss"

### Material and methods

#### Cofilin-actin rods immunohistochemistry

Free-floating 30 μm-thick coronal mice (AβPPswe/PS1A246E, Alzheimer's disease mouse model) brain sections, obtained with the cryostat, were permeabilized with -20ºC methanol, for 1 minute, and then change to fresh -20 ºC methanol for another 5 minutes. Next, tissues were rinsed 3x with PBS, and then blocked with 10% donkey serum in PBS containing 1% BSA, for 1 hour at RT. This was followed by incubation with primary rabbit anti-cofilin antibody (D3F9; 1:2000; Cell Signaling), diluted in 1% BSA in PBS, ON at 4ºC. On the next day, tissues were washed with PBS (6x, 10 minutes) and incubated with Alexa Fluor-568 goat anti-rabbit IgG antibody (A11011; 1:500; Invitrogen), in 1% BSA in PBS, for 2 hours at RT. Then tissues were washed with PBS (6x, 10 minutes) and incubated with DAPI in PBS (1:100; Bio-Rad) for 15 minutes at RT. After, tissues were washed in 70% ethanol for 5 minutes, and after removing the ethanol, with 0,1% Sudan Black in 70% ethanol for 10 minutes. Finally, tissues were washed with 70% ethanol for 20-30 seconds (2x), and then once with PBS for 15 seconds. All of the previous steps were performed with agitation. The brain sections were then mounted on normafrost slides silane pre-coated (Normax), with ibidi mounting media (without DAPI). The tissues were visualized and photographed using a Leica DMI6000 FFW microscope equipped with a Hamamatsu FLASH 4.0 camera (Hamamatsu, Japan), and the LAS X software, and pictures were taken using a 20x magnification. Number of cofilin-actin rods per area was counted using Fiji software [1].

#### Proteins and peptides

Human myc-tagged (TRTRPL**EQKLISEEDL**AANDILDYKDDDDKV) S100B was expressed in *E. coli* (BL21 (DE3) E. Cline Express, Lucigen) and purified to homogeneity as described [42,43]. S100B-myc concentrations were estimated as homodimer equivalents using by UV spectroscopy at 280 nm using the theoretical extinction coefficient value of $\varepsilon_{280\ nm}$ = 5,960 $M^{-1}cm^{-1}$

#### S100B internalization by neurons

Primary neurons (DIV 7) were treated during 30 minutes with S100B myc tag (2.5 μM) and then fixed with 4% PFA in 1x cytoskeleton preservation buffer (10mM MES pH 6.1; 138mM KCl; 3mM $MgCl_2$; 2mM EGTA pH 7; 0.32M sucrose), for 30 minutes at RT. After fixation, cells were washed with PBS (3x, 5 minutes) and permeabilized with 0,1% Triton X-100 in PBS for 10 minutes at RT. Cells were, once again, washed with PBS (3x, 5 minutes) and blocked with 5% BSA in PBS for 1 hour, then incubated with primary mouse MYC tag 4A6 antibody (05-724; 1:500; Merck Millipore) and rabbit βIII tubulin (302302; 1:1000; Synaptic Systems) in 1% BSA in PBS, ON at 4 ºC. On the next day, cells were washed with PBS (3x, 5 minutes) and incubated with Alexa Fluor-488 goat anti-mouse IgG antibody (A11029;1:1000;

Invitrogen) and Alexa Fluor-568 goat anti-rabbit IgG antibody (A11011; 1:1500; Invitrogen) with 1% BSA in PBS for 1 hour at RT, and then washed once again with PBS (3x, 5 minutes) After this, coverslips were mounted with FluoroshieldTM with DAPI (Sigma-Aldrich). Cell visualization and image capture were done using the Zeiss Axio Imager Z1 microscope equipped with an Axiocam MR3.0 camera and Axivision 4.9.1 software, with a 40x magnification.

**S100B expression by neurons**

Primary neurons (DIV 7) were fixed with 4% PFA in 1x cytoskeleton preservation buffer (10mM MES pH 6.1; 138mM KCl; 3mM $MgCl_2$; 2mM EGTA pH 7; 0.32M sucrose), for 30 minutes at RT. After fixation, cells were rinsed 3x with PBS, and then permeabilized with 0,3% Triton X-100 in PBS for 15 minutes, at RT. Next, cells were rinsed 3x with PBS again, and blocked with BSA 5% in PBS for 1 hour, at RT. After blocking, neurons were incubated with primary mouse anti-βIII tubulin antibody (1:2000; Promega) and with primary rabbit anti-S100B antibody (1:200; Abcam), in BSA 1% in PBS, ON at 4 ºC. On the next day, cells were rinsed 3x with PBS, and incubated with Alexa Fluor-488 goat anti-mouse IgG antibody (1:1500; Invitrogen) and Alexa Fluor-568 goat anti-rabbit IgG antibody (1:1500; Invitrogen) at 1% BSA in PBS for 1 hour at RT. After this, cells were rinsed 3x with PBS and coverslips were mounted with FluoroshieldTM with DAPI (Sigma-Aldrich). Cell visualization and image capture was done using the Zeiss Axio Imager Z1 microscope equipped with an Axiocam MR3.0 camera and Axivision 4.9.1 software, with a 40x magnification.

**Cofilin and βIII tubulin immunocytochemistry in primary rat neurons**

DIV5 rat neurons were incubated with 10 μM Aβ42 oligomers for 24 or 48 hours or at DIV6 with 1.0, 2.5, 5.0, 10.0 μM S100B for 24 hours; S100B was added halfway through the 48-hour Aβ42 treatment. Neurons were fixed at DIV 7 with 4% PFA in 1x cytoskeleton preservation buffer (10mM MES pH 6.1; 138mM KCl; 3mM $MgCl_2$; 2mM EGTA pH 7; 0.32M sucrose), for 30 minutes at RT, and then rinsed 3x times with PBS, and incubated with -20ºC methanol, for 3 minutes at RT. Next, cells were again rinsed 3x with PBS and incubated with blocking solution (2,5% donkey serum in PBS containing 1% BSA (Bovine serum albumin), for 1 hour at RT.  This was followed by incubation with primary rabbit anti-cofilin antibody (D3F9; 1:2000; Cell Signaling) and mouse anti-βIII tubulin antibody (G712A; 1:2000; Promega), diluted in 1% BSA in PBS, overnight (ON) at 4ºC. On the next day, neurons were rinsed 3x with PBS and incubated with Alexa Fluor-568 goat anti-rabbit IgG antibody (A11011; 1:1000; Invitrogen) and Alexa Fluor-488 goat anti-mouse IgG antibody (A11029;1:1000; Invitrogen), in 1% BSA in PBS, for 1 hour at RT. After incubation, neurons were rinsed 3x with PBS and coverslips were mounted with FluoroshieldTM with DAPI (Sigma-Aldrich). Cell visualization and image capture was done using the Zeiss Axio Imager Z1 microscope equipped with an Axiocam MR3.0 camera and Axivision 4.9.1 software, with a 40x magnification. Finally, cells with cofilin-actin rods were counted and divided per total number of cells.

#### Results

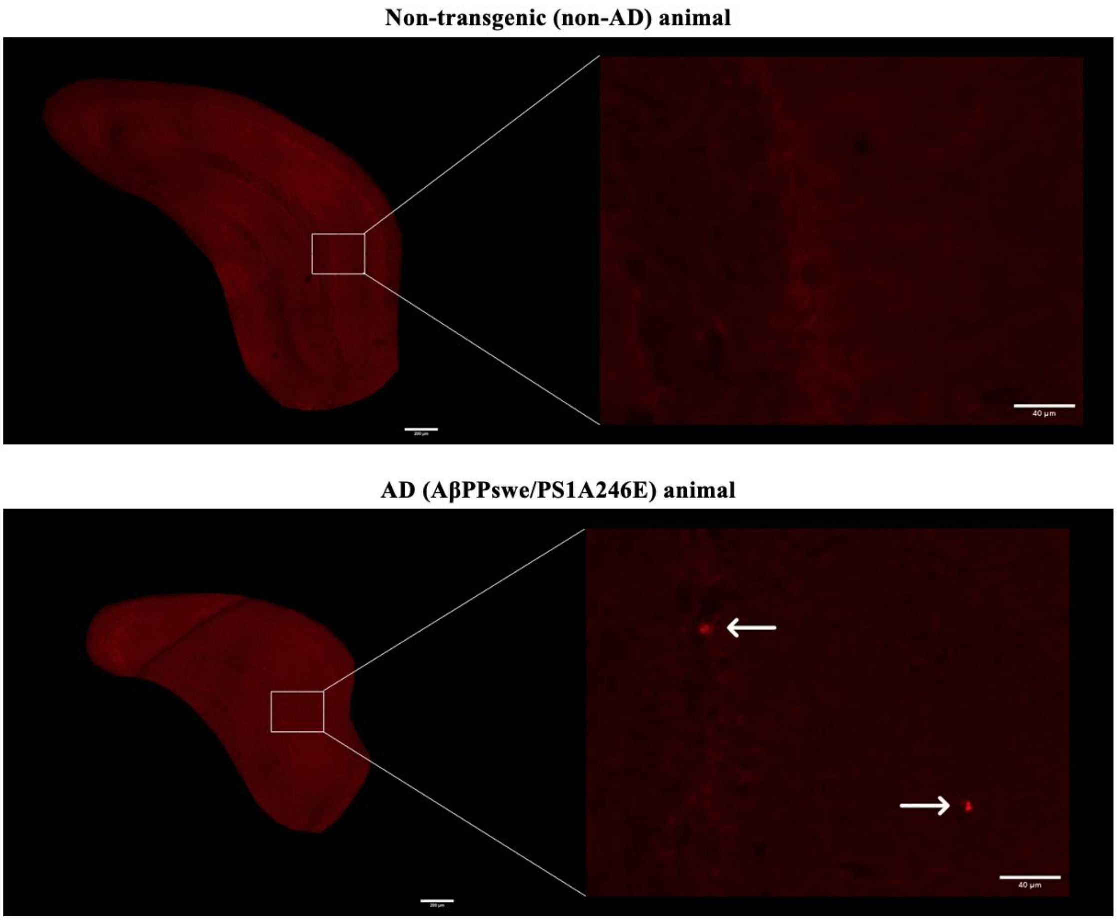


**Figure S1. Cofilin-actin rod formation in hippocampus of NT and AD mice.** Cofilin-actin rod formation (arrows) in the hippocampus of AD (AβPPswe/PS1A246E) and non-transgenic (non-AD) animals, at 7 months. Red: Cofilin. Hippocampus Scale bar = 200 μm; Enlarged insertion, scale bar = 40 μm.

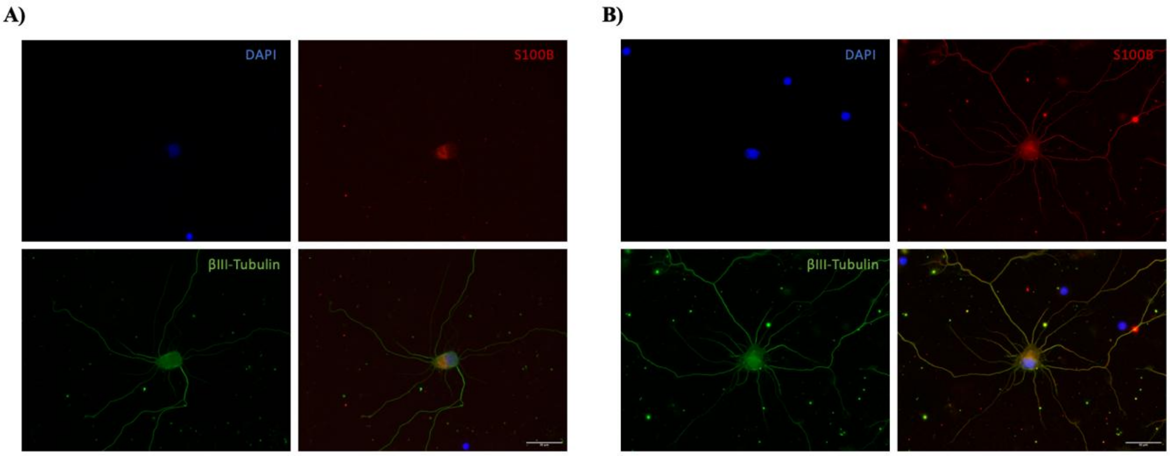


**Figure S2.** S100B expression and internalization in primary cultures of rat hippocampal neurons. Endogenous expression **(A)** and exogenous internalization **(B)** of S100B in rat hippocampal neurons (DIV7). Green: βIII-Tubulin; Red: S100B. Blue: DAPI. Scale bar = 30 μm.

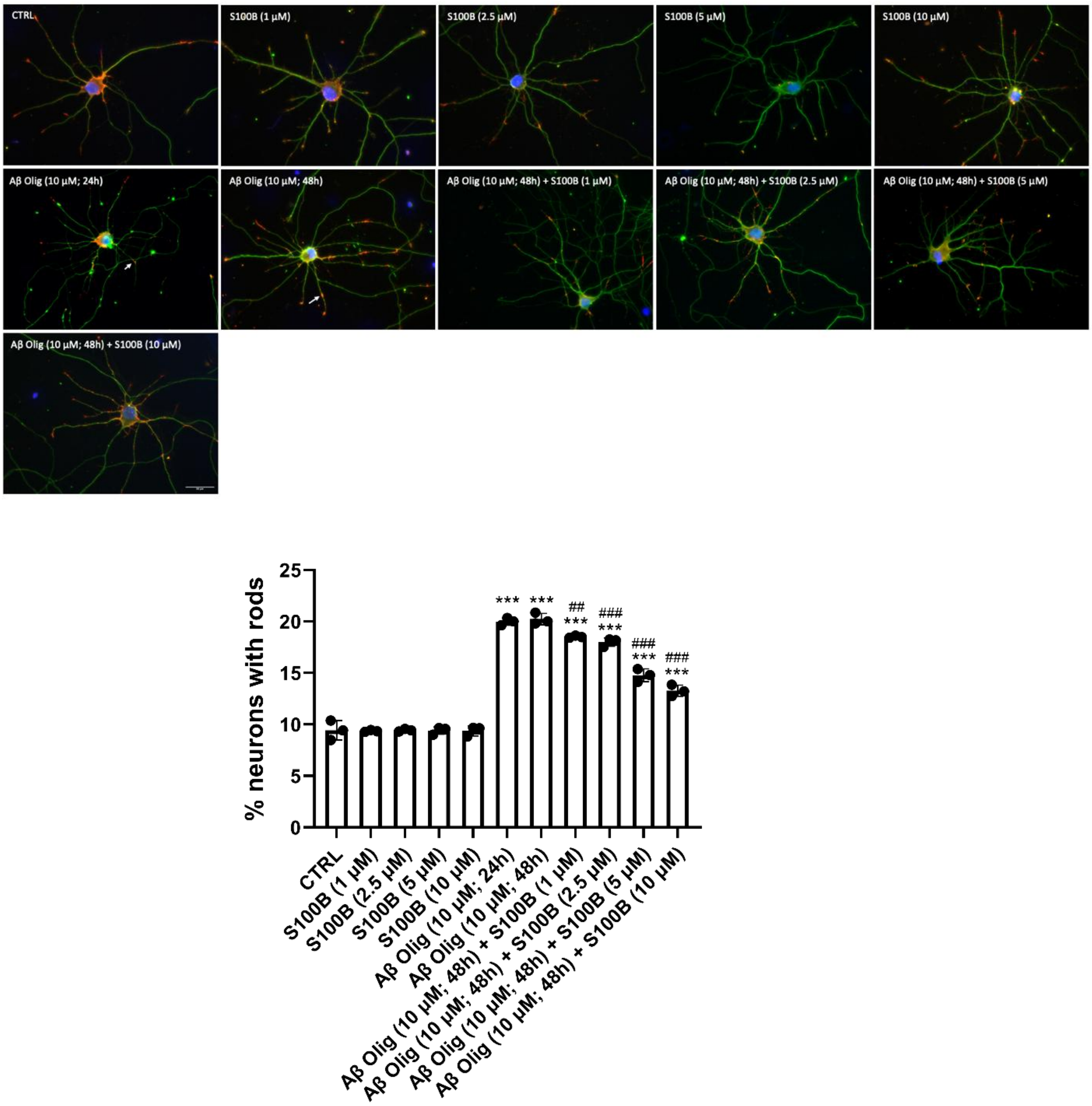


**Figure S3. Impact of S100B on Aβ oligomers-induced cofilin-actin rod formation in primary cultures of rat hippocampal neurons.** Representative images and quantification plots of rat hippocampal neurons (DIV7) showing increased % of cells with cofilin-actin rods after incubation with Aβ oligomers (Aβ Olig) 24h or 48h (10 µM). S100B (1; 2.5; 5; 10 µM) added after 24 hours partially rescued the Aβ-induced phenotype in a dose dependent manner. S100B alone did not produce alterations, as compared to control (CTRL). White arrows point out cofilin-actin rods. Green: βIII-Tubulin; Red: Cofilin. Blue: DAPI. Scale bar = 30 µm. Data are expressed as mean ± SD. (n= 3 per group). * or $^{\#}$ $p<0.05$; *** or $^{\#\#\#}$ $p<0.001$. (* comparation with control; $^{\#}$ comparation with Aβ oligomers 48h).
